# Lysosomal accumulation of Masitinib alters autophagy via pH-dependent trapping

**DOI:** 10.64898/2025.12.07.692822

**Authors:** Abdulrahman El Sayed, Monika Kluzek, Remigiusz Serwa, Abdelhalim Azzi

## Abstract

Masitinib is an oral tyrosine kinase inhibitor developed initially for the treatment of mastocytosis. Preclinical studies suggest that masitinib can modulate mast cell activity and neuroinflammation, making it a potential therapeutic candidate for neurodegenerative diseases. However, clinical outcomes remain inconclusive, which may be linked to a lack of understanding of its molecular mechanism of action and potential off-target effects. Using a panel of cancer cell lines, we found that masitinib suppresses mTORC1 activity while simultaneously inducing AKT phosphorylation. Interestingly, thermal proteome profiling analysis revealed that lysosomal proteins were the most thermally affected by masitinib. Using large unilamellar vesicle systems that mimic the lipid composition and physicochemical environment of lysosomal membranes, we observed that masitinib preferentially accumulates in acidic vesicle membranes, consistent with a pH-dependent trapping mechanism. Furthermore, cellular assays demonstrated that masitinib disrupts autophagy by impairing lysosomal acidification. Together, these findings elucidate the lysosomal accumulation of masitinib, providing a molecular basis for its mechanism of action.

## Introduction

Masitinib is an orally administered small-molecule tyrosine kinase inhibitor (TKI) that targets activated cells of the innate neuroimmune system. In vitro, masitinib is able to inhibit the activity of several receptor and non-receptor tyrosine kinases, including c-Kit, Lyn, PDGFR-α, PDGFR-β, Abl1, c-Fms, and Src, which are critically involved in immune cell activation, inflammatory signaling, and oncogenic proliferation [1–2]. Recent preclinical studies have highlighted the neuroprotective potential of masitinib. Indeed, in an acrolein-induced Alzheimer’s disease (AD) mouse model, masitinib attenuated disease pathology by reducing phosphorylated tau levels and suppressing neuroinflammation through the inhibition of the NF-κB/NLRP3/caspase-1 inflammatory pathway and microglial activation [3]. Similarly, in a model of experimental autoimmune encephalomyelitis (EAE), masitinib mitigated neuronal damage by reducing pro-inflammatory cytokines [4]. Together, these findings support the hypothesis that modulation of innate immune responses underlies the beneficial effects of masitinib in neurodegenerative and neuroinflammatory disorders. However, despite these promising preclinical results, the clinical translation of masitinib remains challenging. In fact, a recent meta-analysis of randomized controlled trials demonstrated a significantly higher incidence of adverse events in patients receiving masitinib compared with controls, regardless of adverse event grade or dose [5]. Moreover, a phase 3 clinical study in AD patients reported inconclusive efficacy for masitinib at 6.0 mg/kg/day compared to placebo (n = 186 and 91, respectively) [6]. These observations highlight the need for further investigation into the exact molecular mechanism of action of this drug. Indeed, at the molecular level, the intracellular mechanisms underlying the pharmacological effects of masitinib remain poorly described. Although an in vitro assay study demonstrated that the primary targets of masitinib are receptor tyrosine kinases (RTKs) [1], cellular studies investigating its molecular mechanism of action are still lacking. For example, the downstream signaling consequences of masitinib on the activity of the PI3K/Akt/mTOR and MAPK pathways, which are known to be directly under the control of RTK, are poorly understood [7–8]. These signaling pathways are central regulators of cell growth, survival, and metabolism. Moreover, it is very well established that several RTKs are often dysregulated in cancer. Therefore, understanding how masitinib affects the activity of these signalling pathways in various cellular models may shed light on both its therapeutic potential and off-target effects. For instance, in pancreatic cancer cell lines, masitinib alone exhibited limited antiproliferative activity; however, when combined with gemcitabine, it showed synergistic effects, particularly in gemcitabine-refractory models [9]. Interestingly, a study using c-Kit-negative hepatocellular carcinoma (HepG2) cells revealed that masitinib induces reactive oxygen species dependent apoptosis, suggesting additional mechanisms independent of c-Kit inhibition [10]. Besides the lack of understanding of its mechanism of action, masitinib might also accumulate in lysosomes. Indeed, studies have shown that several receptor tyrosine kinase inhibitors, such as imatinib, sunitinib, and lapatinib, undergo lysosomal sequestration via protonation within acidic compartments, a process that can alter intracellular distribution, pharmacokinetics, and target engagement [11–12]. Lysosomal trapping not only reduces drug availability in the cytosol but also impairs lysosomal function or induces stress responses [13]. In this respect, Emilia Neuwirt et al. [14] have shown that masitinib activates the NLRP3 inflammasome and induces lysosome swelling. Whether these alterations result from the direct activation of the inflammasome or are linked to the accumulation of the drug in the lysosome is currently unknown.

To better understand this phenomenon, we first used thermal proteome profiling. Our data indicate that lysosomal function is a significant off-target effect of masitinib. In fact, a large number of proteins that showed thermal stability changes upon treatment with masitinib localize to the lysosome. Next, to examine whether masitinib could preferentially accumulate in the lysosome, we used lipid model systems to mimic both the lipid composition and the physicochemical environment of the acidic intracellular compartments [15]. To that end, we monitored shifts in the fluorescence spectra of two lipophilic dyes sensitive to lipid packing, namely Laurdan [16] and Di-4-ANEDPPHQ [17], which would reflect insertion of masitinib within the membrane.

Using this biophysical approach, we observed that masitinib preferentially accumulates within the membrane of acidic vesicles, supporting the hypothesis of pH-dependent trapping. Moreover, our data show that masitinib enhances autophagosome formation while its accumulation in lysosomes disrupts lysosomal acidification, leading to an alteration in autophagic flux. This experimental system provides mechanistic insight into the lysosomal accumulation of masitinib and helps interpret its intracellular localization and functional consequences observed in cellular models.

## Results

### Masitinib enhances AKT S473 phosphorylation

The protein kinase B (PKB), also known as Akt, functions as a key signaling protein downstream of receptor tyrosine kinases (RTKs), and its phosphorylation level provides a functional readout of RTK pathway activation. Changes in AKT S473 phosphorylation reflect not only the canonical PI3K/mTORC, but also non-canonical input proteins such as DNA-PK, TBK1, and IKKα [18]. Thus, this phosphorylation site of AKT is a highly informative marker for both primary pathway activity and feedback responses to RTK inhibition. Using a variety of cancer cell lines, we examined the impact of masitinib on AKT S473 phosphorylation. This experiment demonstrated that masitinib treatment resulted in a substantial increase in AKT1 phosphorylation in several leukemia cell lines and the breast cancer cell line MCF7 (Figure 1A**, Supplementary Figure 1**). In stark contrast with AKT phosphorylation, our data show that masitinib alters cell growth in these cell lines. Indeed, increasing concentrations of masitinib result in a marked decrease in cell growth (**Figure 1B**). Both changes in AKT phosphorylation and cell growth were robust and reproducible across independent experiments. Similar results were observed in several genetically and phenotypically distinct cell lines, indicating a broad cellular response to the drug rather than a cell-line-restricted artifact.

**Figure 1.**
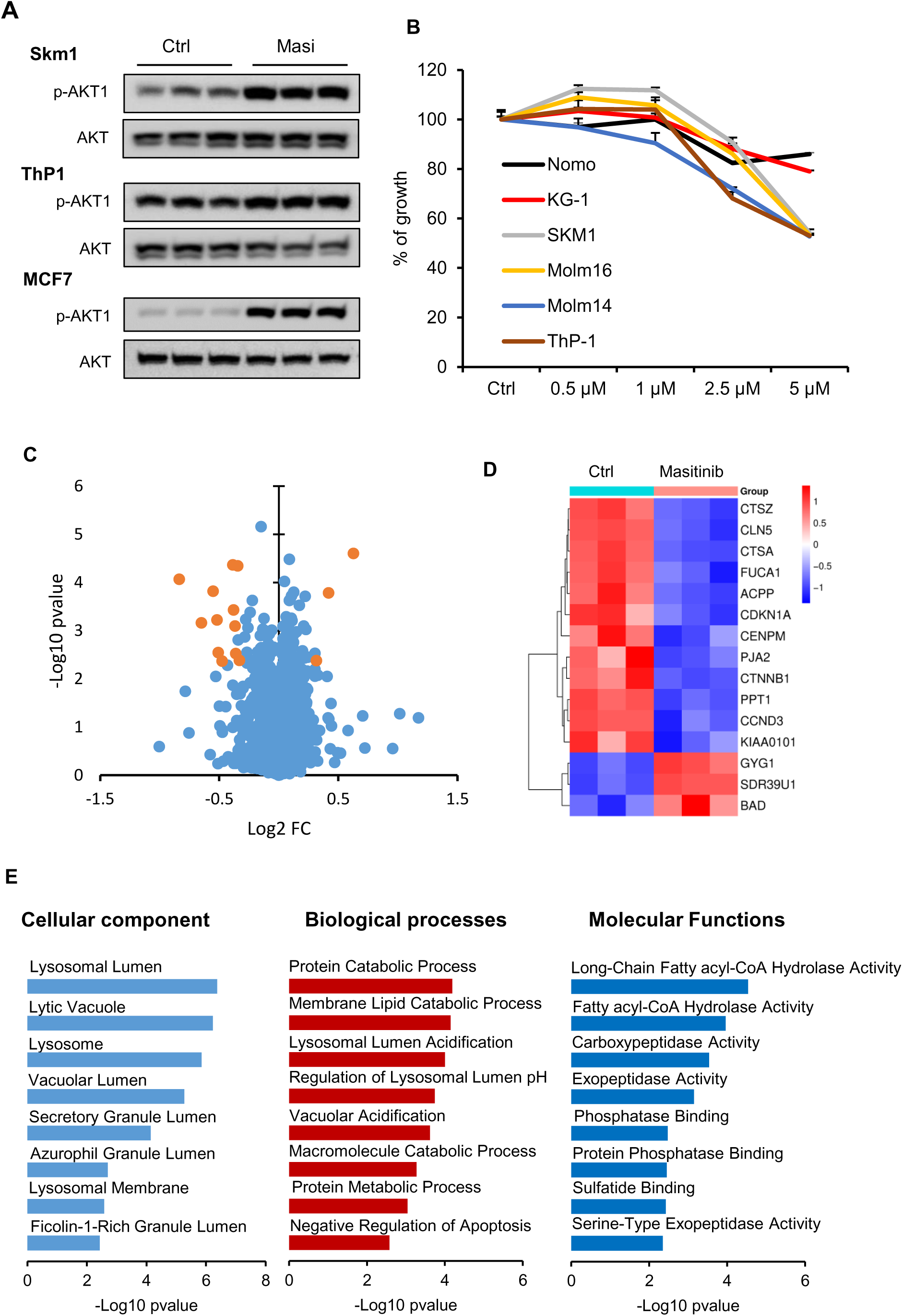
Masitinib alters the AKT1 phosphorylation. (**A**) Lysate from Skm1, ThP-1, and MCF7 cells control or treated with 5 µM Masitinib (Masi) for 24hours, then stained for the indicated antibodies. N = 3 separate biological replicates per condition. (**B**) Line graph showing the percentage (with respect to control [Ctrl], 100 %) of viable cells after treatment with the indicated concentration of masitinib for 72 h. The data are presented as the means ± SEMs (n = 6). One-way ANOVA with multiple comparisons: Dunnett’s test. (**C**) Proteome Integral Solubility Alteration Analysis of Masitinib. Volcano plot of PISA results expressed as log-transformed values of the ratio of protein soluble amount per protein of Masitinib-treated compared to control-treated cells, and relative-log10 p-value. Each dot represents an individual protein identified and quantified, with a minimum of two peptides included in the final analysis. The most significant hits are highlighted in orange font (Log2FC>0.3, –Log10 p-value>2.2). (**D**) Heatmap showing the normalized protein abundance (solubility) of the most significant identified hits. Note the consistent observation between the biological replicates of each condition. Clustering method: average linkage. Distance Measurement Method: Kendalĺs Tau. Raw Z-score: –1.5 to 1.5. (**E**) Enrichment analyses on GO terms: cellular localization (CC), biological processes (BP), and molecular functions of the thermally altered proteins by masitinib. Results of the bar graphs highlight the top affected terms.

### Thermal stability profiling reveals lysosomal proteins as the top thermally altered proteins by masitinib

To screen for potential targets of masitinib, we used the Proteome Integral Solubility Alteration (PISA) assay. This experimental approach enables the assessment of drug-induced alterations in protein thermal stability across the entire cellular proteome [19–21]. SKM-1 cells were treated with 20 µM masitinib or vehicle control for 2 hours. Cells were subjected to a defined temperature gradient; soluble protein fractions were collected, digested with trypsin, TMT-labeled, and analyzed by quantitative LC-MS/MS. In total, 5953 protein groups were identified with high confidence across control and treated cells (**Supplementary Table 1)**. This analysis showed minimal changes in the thermal stability of the proteome. Compared to vehicle-treated cells, only 15 proteins showed significant changes in thermal stability, of which 12 showed lower, and 3 showed higher thermal stability upon masitinib treatment (p-value <0.01, Log2FC >0.3) (**Figure 1 C-D, Supplementary Table 1**). Interestingly, among the most significantly altered proteins, we observed a prominent cluster of lysosomal enzymes and lysosome-associated proteins, including PPT1 (palmitoyl-protein thioesterase 1), CTSZ (cathepsin Z), CTSA (cathepsin A), CLN5 (neuronal ceroid lipofuscinosis protein 5), and FUCA1 (α-L-fucosidase 1) [22] (**Figure 1 D**). In line with the functions of these enzymes, pathway analysis revealed that thermally affected proteins by masitinib localize mainly to the lysosome (**Figure 1 E**), and they play a key role in lysosomal proteolysis, processing of glycoproteins, or lipid/protein turnover (**Figure 1 E**). These observations suggest that masitinib may perturb lysosomal integrity or enzymatic activity. Moreover, the presence of several functionally related lysosomal enzymes argues against random protein destabilization. Instead, it suggests either (I) a direct interaction of masitinib with lysosomal proteins or (II) the accumulation of masitinib in lysosomes, which in turn alters their function.

In addition to lysosomal proteins, masitinib alters the thermal stability of several proteins involved in cell-cycle and DNA replication, such as KIAA0101, a PCNA-binding factor linked to DNA replication stress and repair; CCND3 (Cyclin D3), which regulates G1/S transition; CDKN1A/p21, a cyclin-dependent kinase inhibitor; and CENPM, a centromere-associated protein involved in centromere assembly [22] (**Figure 1 C-D**). These changes suggest that masitinib may affect the cell cycle either directly via its RTK target or through a separate mechanism. In summary, the thermal proteome profiling data for masitinib consistently indicate that lysosomal function is a major off-target effect of this inhibitor, which is accompanied by secondary effects on cell-cycle regulators. The lysosomal signature provided the basis for subsequent investigations of the upstream factors driving changes in lysosomal protein alterations and the downstream consequences involving autophagy and AKT phosphorylation.

### Masitinib mediated AKT S473 phosphorylation is independent of class I PI3K but requires VPS34 activity and is rapamycin-sensitive

Our data above showed that masitinib leads to an increase in AKT phosphorylation and disrupts the thermal stability of several lysosomal enzymes, suggesting a functional interplay between lysosome integrity and stress-mediated AKT activation. To study this connection, we examined whether canonical PI3K and mTOR pathways mediate the paradoxical induction of AKT phosphorylation upon RTK inhibition by masitinib.

We first asked whether class I PI3 Kinase activity is required for masitinib mediated AKT phosphorylation. We used GDC-0980 and wortmannin, two well-known inhibitors of class I PI3K [23–24]. When used alone, both inhibitors led to a decrease in AKT phosphorylation. However, when combined with masitinib, the increase of AKT S473 phosphorylation was still preserved (**Figure 2 A-B, Supplementary Figure 2 A**), indicating that the increase in AKT phosphorylation upon masitinib treatment does not depend on classical PI3K/PIP3/PDK1 signaling [18], and suggests the involvement of a separate upstream pathway. We next examined the role of mTORC1/mTORC2 signaling using rapamycin. Interestingly, simultaneous treatment with rapamycin suppresses the increase in AKT S473 phosphorylation induced by masitinib, suggesting that the AKT activation mediated by masitinib requires mTOR signaling (**Figure 2 A-B, Supplementary Figure 2A**). Short-term treatment (6 hours) with rapamycin primarily inhibits mTORC1, whereas prolonged exposure also suppresses mTORC2 activity [25]. Moreover, masitinib treatment led to a significant decrease in phosphorylated P70S6 kinase at Thr389, a known direct target of mTORC1 (**Supplementary Figure 3 A-C**), suggesting that masitinib mediated AKT phosphorylation could be linked to mTORC2. To examine the contribution of mTORC2 to masitinib-mediated AKT phosphorylation, we used MCF7 cells stably expressing either a control shRNA or shRNAs targeting Rictor, a key component required for mTORC2 function [26]. As shown in **Supplementary Figure 3D-E**, Rictor depletion did not completely block masitinib-mediated AKT phosphorylation, but significantly reduced it, suggesting that mTORC2 plays a major, but not absolute, role in mediating this effect.

**Figure 2.**
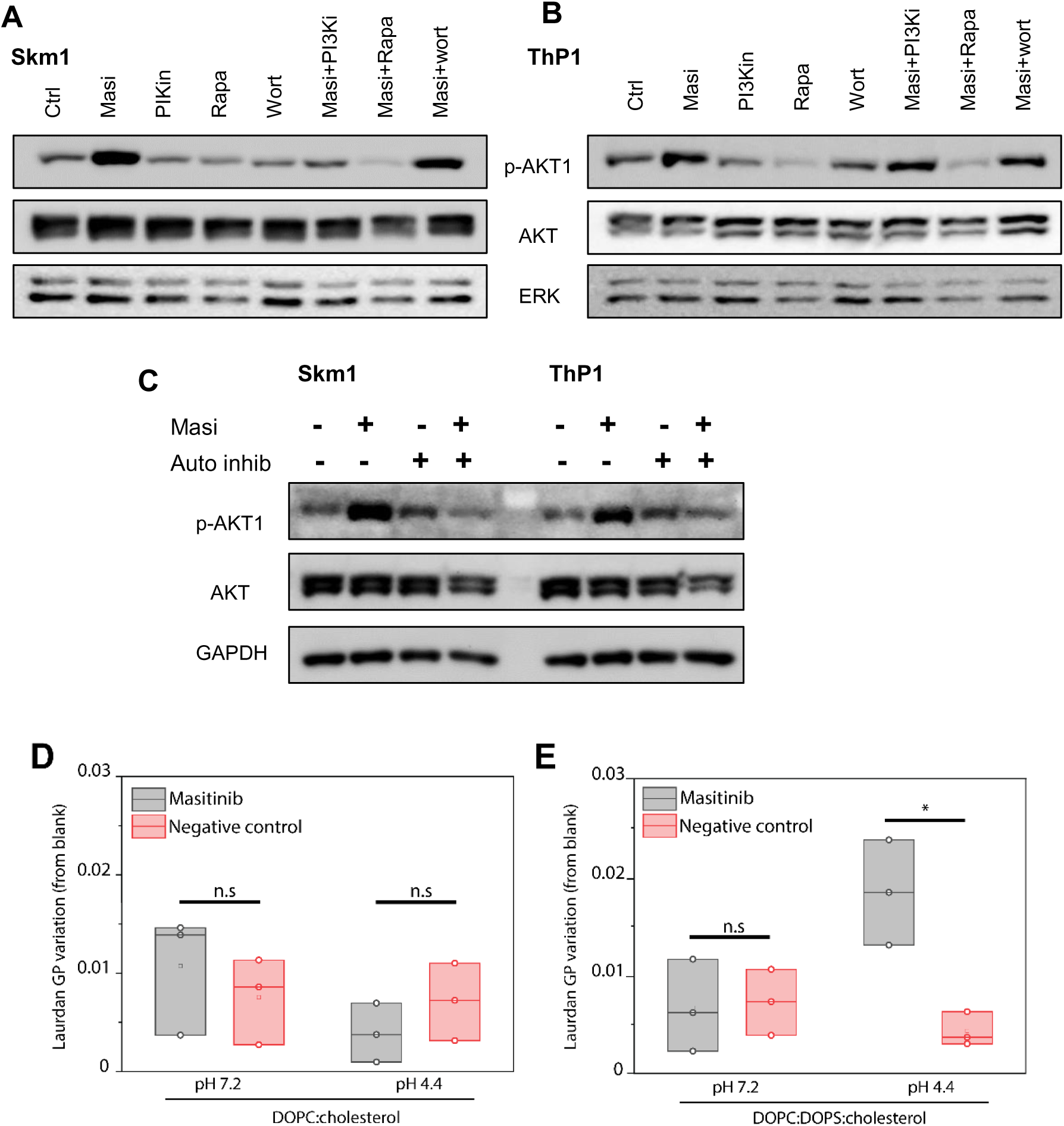
Masitinib-mediated AKT S473 phosphorylation is independent of class I PI3K. (**A-B**) Lysate from Skm1, ThP-1 cells control or treated with 5 µM Masitinib (Masi) supplemented with the indicated inhibitor for 24 hours, then stained for the indicated antibodies. PI3Ki (GDC-0980, 50nM), Rapamycin 50nM, wortmannin 50nM. (**C**) Lysate from Skm1, ThP-1 cells control or treated with 5 µM Masitinib (Masi) supplemented or not with 200 nM VPS34 inhibitor for 24 hours, then stained for the indicated antibodies. (**D-E**) Masitinib possesses high affinity towards membrane lipids. Variation of Laurdan GP for either DOPC: cholesterol (**D**) or DOPC: DOPS: cholesterol (**E**) LUVs incubated with either 5 μM Masitinib (black) or DMSO only (negative control, red) at either pH 7.2 or 4.4. Corrected GP values were obtained by subtracting GP values of LUVs alone (blank). A positive increase in GP indicates tighter lipid packing in the membrane due to Masitinib insertion. Each data point represents one technical repeat. Statistical analysis was performed using one-way ANOVA, with a confidence value of 0.05. * – p < 0.05.

Given its established role in autophagy initiation and PI3P generation [27], we next examined whether class III PI3K (VPS34) activity contributes to masitinib mediated AKT phosphorylation. As shown in **Figure 2C**, pharmacologic inhibition of VPS34 also prevented AKT S473 induction, implicating VPS34 as a necessary component of this signaling axis.

Taken together, our data demonstrate that the increase in AKT phosphorylation following masitinib treatment does not require canonical class I PI3K activity, but instead depends on VPS34 and is sensitive to rapamycin. Together with thermal profiling results, these findings suggest that masitinib might accumulate within lysosomal compartments and alter its function, which in turn triggers a compensatory activation of AKT through a non-canonical pathway.

### Masitinib preferentially interacts with membranes mimicking the lysosomal milieu

Following our observation indicating a substantial change in the thermal stability of lysosomal proteins upon masitinib treatment (**Figure 1**), we wanted to assess whether Masitinib preferentially interacts with lysosomal membranes and if the lysosomal milieu (namely acidic pH and lysosomal membrane lipid composition) plays a role in such interaction.

To determine whether Masitinib integrates into lipid membranes, we evaluated changes in the fluorescence emission spectra of Laurdan, a lipophilic dye. Laurdan’s emission spectrum shifts based on the polarity of its environment, alternating between two primary peaks at 440 nm and 490 nm. In lipid membranes, these spectral changes reflect lipid packing in the bilayer: tightly packed membranes predominantly show the 440 nm peak, while loosely packed membranes favour the 490 nm peak [16, 28].

Lipid packing information is described by the Laurdan General Polarization (Laurdan GP) [16], which indicates the relative intensity of the two spectral peaks. A high GP (>0) signifies densely packed membranes, whereas a low GP (<0) indicates lower lipid surface density. Changes in Laurdan GP under specific conditions can reveal alterations in membrane packing. For instance, a Masitinib inserting into the membrane surface is expected to increase GP, as lipids pack more tightly to accommodate the Masitinib [29]. Large unilamellar vesicles (LUVs) were prepared to mimic the lipid composition of either the early endosomal membrane [30], (DOPC: cholesterol, 60:40 mol/mol, PBS, pH 7.2, 297 mOsm/Kg) or the inner leaflet of the lysosomal membrane, enriched in negatively charged lipids [31], as to assess whether Masitinib insertion in the membrane would occur preferentially at different stages of the endocytic pathway. For control, we measured changes in Laurdan spectra of LUVs in buffer only (blank) and LUVs incubated with DMSO (negative control).

In the kinetic profile for both lipid compositions and pHs (**Supplementary Figure 4**), we observe a characteristic drop in Laurdan GP within 60 min of incubation for both Masitinib and negative control samples; such an effect is due to sample equilibration to 37°C as LUVs membrane becomes less packed due to the temperature increase. Kinetic profiles of DOPC: cholesterol LUVs showed no insertion of Masitinib at either pH 7.2 or 4.4, as the kinetic profiles for Masitinib or negative control show no difference in Laurdan GP throughout the incubation period (**Supplementary Figure 4**). The addition of 20% negative charge was also not sufficient to affect Masitinib insertion at pH 7.2.

However, the presence of a 20% negative charge by the addition of DOPS at pH 4.4, led to a clear change in Laurdan GP between samples incubated with Masitinib and the negative control. Particularly, we observe that LUVs incubated with Masitinib (black line) display a higher Laurdan GP than the negative control (red line), and that such a higher GP value remains constant throughout the 8 h incubation period, suggesting a stable insertion within the membrane (**Supplementary Figure 4**).

Analysis of corrected GP values revealed that DOPC: cholesterol LUVs at both pH 7.2 and 4.4 exhibited no significant difference between Masitinib and negative control samples (**Figure 2D**). This suggests that the observed GP variations result from DMSO’s known effect of dehydrating lipid bilayers, leading to tighter packing [32]. Similarly, DOPC: DOPS: cholesterol LUVs at pH 7.2 showed a minor GP increase but no statistical difference between Masitinib and negative control (**Figure 2E**). However, for DOPC: DOPS: cholesterol LUVs at pH 4.4, a statistically significant difference was observed, with the negative control causing a negligible GP increase, while Masitinib raised Laurdan GP by ∼0.02 (**Figure 2E**).

To further validate that the lysosomal milieu is driving the membrane-Masitinib interaction, we probed changes in Laurdan spectra in membranes that more closely resemble the biological complexity of real lysosomal membranes. To that end, we performed the Laurdan kinetics measurements we conducted in simpler LUVs compositions (**Figure 3**) in liposomes composed of POPC: DOPE: Sphingomyelin (SM): Bis(monoacylglycerol) phosphate (BMP): cholesterol 25:5:30:15:25 (mol/mol %), mimicking the lipid compositional complexity of lysosomes [33], at either pH 7.2 or pH 4.4.

**Figure 3:**
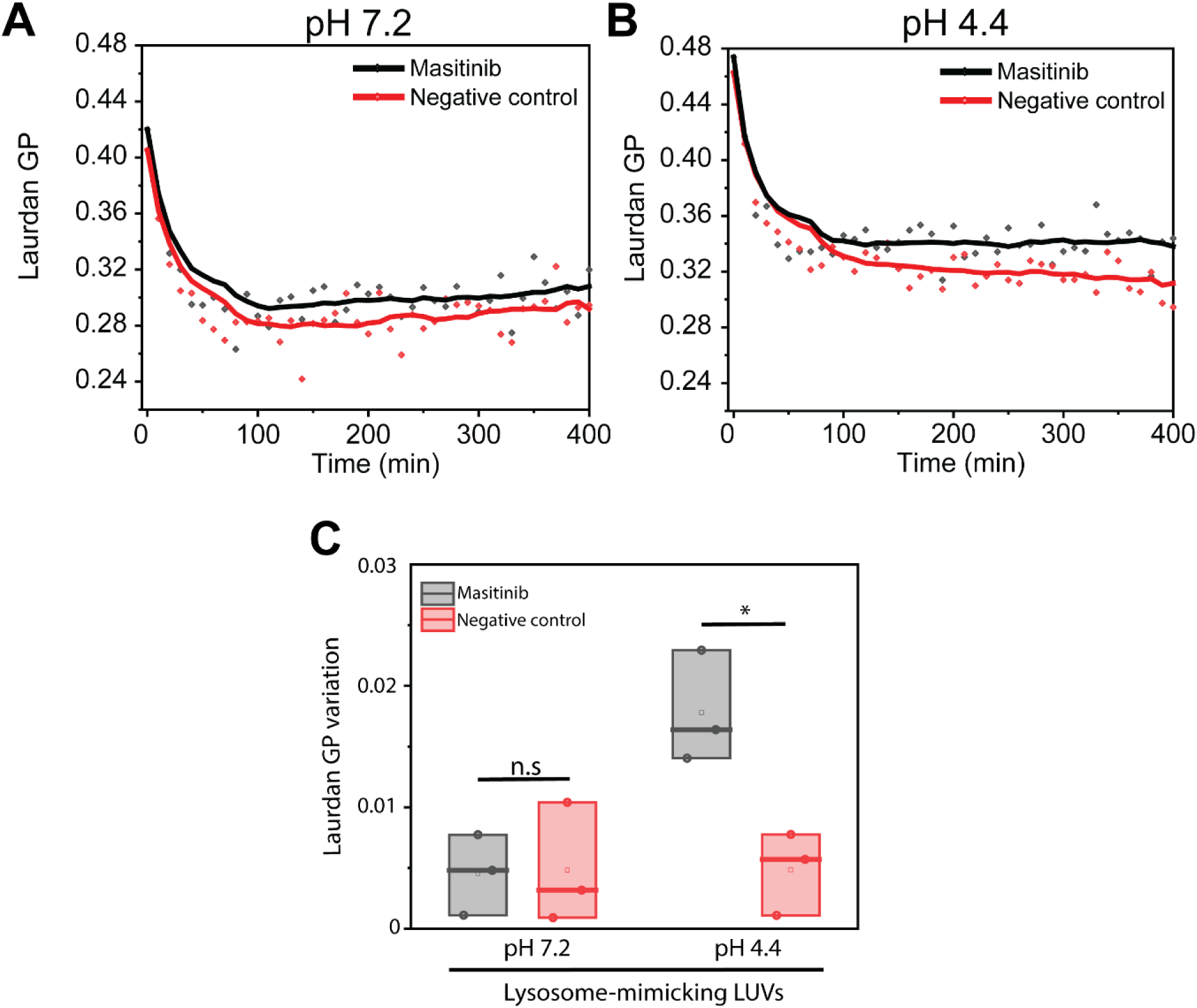
Representative kinetics profile of Laurdan GP for liposomes composed of POPC: DOPS: SM: BMP: PE: cholesterol (mimicking the lipid complexity of lysosomal membranes) at pH 7.2. (**A**) and pH 4.4 (**B**), incubated with 5 μM Masitinib in DMSO (black) or DMSO only (negative control, red). Data symbols represent each GP measurement, while the line represents the kinetic profile obtained from smoothing the scatter plot using a 15-point average. (**C**) Variation of Laurdan GP for LUVs composed of POPC: DOPS: SM: BMP: PE: cholesterol with either 5 μM Masitinib (black) or DMSO only (negative control, red) at either pH 7.2 or 4.4. Corrected GP values were obtained by subtracting GP values of LUVs alone (blank). A positive increase in GP indicates tighter lipid packing in the membrane due to Masitinib insertion. Each data point represents one technical repeat. Statistical analysis was performed using one-way ANOVA, with a confidence value of 0.05. * – p < 0.05.

In the kinetic profile for both pHs (**Figure 3A-B**), we observe the characteristic drop in Laurdan GP within 60 min of incubation for both Masitinib and negative control samples; this effect is due to sample equilibration to 37°C as LUVs membrane becomes less packed due to the temperature increase. Kinetic profiles of lysosome-mimicking LUVs at pH 7.2 showed minimal differences in Laurdan GP throughout the incubation period between Masitinib-treated or DMSO only-treated samples (**Figure 3A**), with less than a ∼0.1 variation observed. Dropping the pH to lysosomal conditions (∼4.4), drastically increases the GP variation between Masitinib-treated compared to DMSO-only (**Figure 3B**), with LUVs incubated with Masitinib (black line) display a GP value almost 0.3 higher than DMSO only samples (red line).

Analysis of blank-corrected GP values (**Figure 3C**) revealed that at pH 7.2, Masitinib displays no significant variation in GP value compared to the DMSO negative control. LUVs incubated with Masitinib (black) at pH 4.4 instead display a clear and statistically relevant increase in Laurdan GP values compared to DMSO negative control (red), indicating an increase in membrane order at such conditions due to the interaction (and most likely insertion) of Masitinib within the membrane. Overall, these results further reinforce that both lipid composition and lysosomal pH are essential for Masitinib interaction and intercalation into membranes, with negatively charged lipids driving the compound affinity. Indeed, for LUVs containing 20% mol DOPS (charge –1 at pH 4.4, total charge –20), we observe a Laurdan GP variation of 0.0177 ± 0.049, and for lysosomal-mimicking LUVs, containing 15% BMP (charge slightly lower than –2 at pH 4.4, total charge slightly lower than –30) we have a comparable variation of Laurdan GP at 0.0177 ± 0.038, most likely by the similar total charge between the two-lipid composition.

These results confirm that Masitinib has a high affinity for lysosomal membranes and that this interaction is driven by a combination of the presence of negatively charged lipids (enriched in lysosomal membranes) and a low pH. Both factors are required for a strong interaction between Masitinib and membranes.

### Acidic pH drives Masitinib’s intercalation within lysosomal-mimicking membranes

To further corroborate the Laurdan kinetics results and visualize the effect of Masitinib on membranes, we performed confocal microscopy coupled with spectral imaging of lipid giant unilamellar vesicles (GUVs) labeled with Di-4-ANEPPDHQ (Di-4). Like Laurdan, Di-4 is a ratiometric membrane dye that responds to changes in lipid packing [17, 28], but it is more sensitive to the incorporation of compounds within the hydrophobic core of the membrane. Similar to Laurdan, Di-4 exhibits a shift in its primary emission peak depending on the lipid packing and order of the membranes. Tightly packed bilayers display an emission peak centered at ∼590 nm, while more disordered (fluid) membranes show a peak at ∼640 nm (**Figure 4A**). This information can be summarized by the ratio of fluorescence intensities at 590 nm and 640 nm (I_590_/I_640_), where a ratio >1 indicates a more ordered membrane and <1 indicates a more fluid membrane.

**Figure 4.**
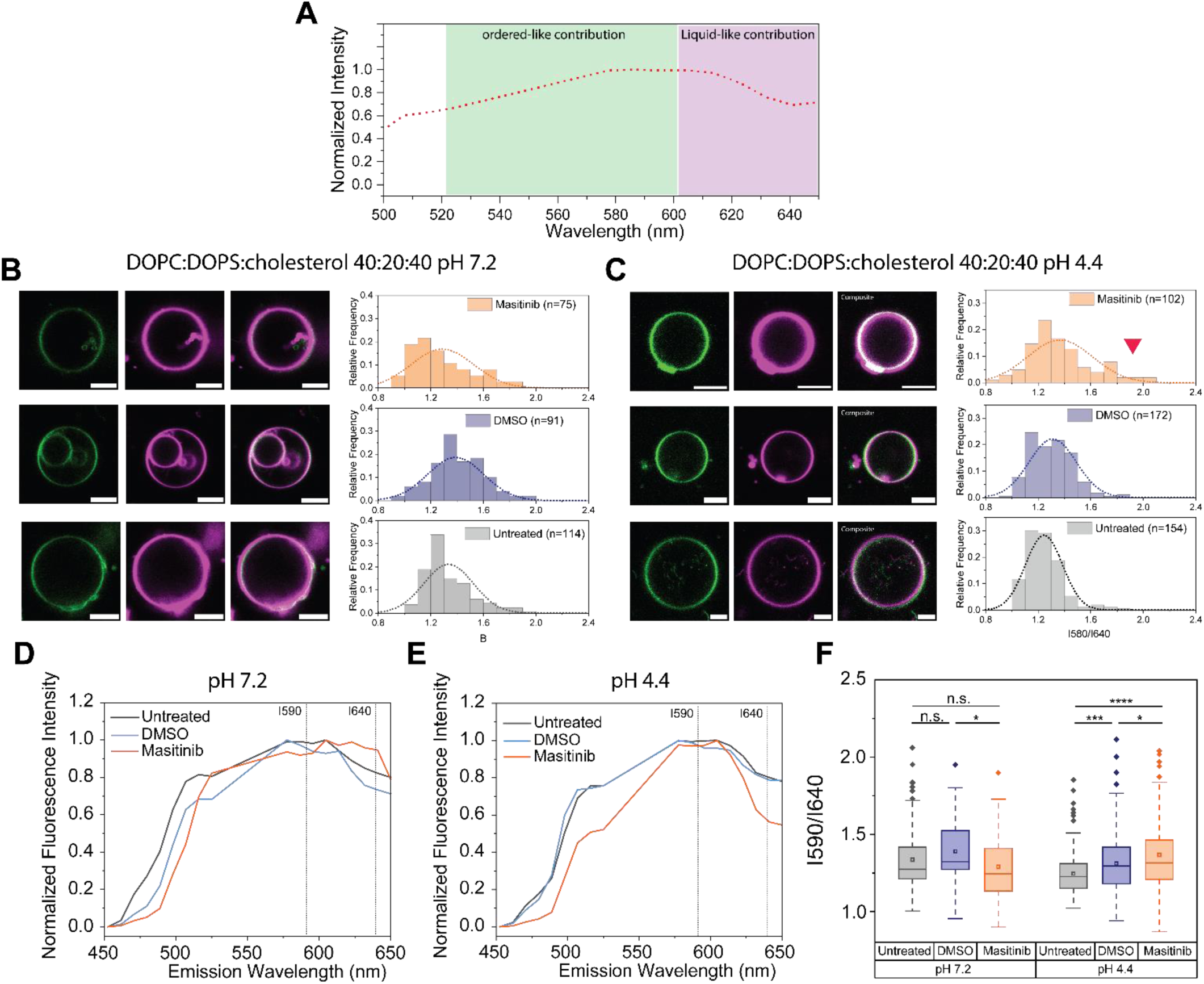
Masitinib intercalates within lysosome-mimicking membranes under acidic conditions. (**A**) Di-4 emission spectra highlighting the two primary contributions: an ordered phase centered at 590 nm (green) and a disordered phase centered at ∼640 nm (magenta). (**B**) Representative confocal images of DOPC: DOPS: cholesterol GUVs labeled with Di-4 at pH 7.2, showing the green (590 nm) and red (640 nm) contributions of Di-4 emission, along with an overlay of both channels for each condition. Corresponding 6I590/I640 ratio distributions were calculated from individual GUVs for untreated (gray), DMSO-treated (blue), and Masitinib-treated (orange) conditions. (**C**) Representative confocal images of DOPC: DOPS: cholesterol GUVs labeled with Di-4 at pH 4.4, showing the green (590 nm) and red (640 nm) contributions of Di-4 emission, along with an overlay of both channels for each condition. Corresponding I590/I640 ratio distributions were calculated from individual GUVs for untreated (gray), DMSO-treated (blue), and Masitinib-treated (orange) conditions. (**D**) Representative Di-4 emission spectra obtained via spectral imaging from DOPC: DOPS: cholesterol GUVs at pH 7.2 for untreated (gray), DMSO-treated (blue), and Masitinib-treated (orange) conditions. (**E**) Representative Di-4 emission spectra obtained via spectral imaging from DOPC: DOPS: cholesterol GUVs at pH 4.4 for untreated (gray), DMSO-treated (blue), and Masitinib-treated (orange) conditions. (**F**) Summary of I590/I640 ratio distributions obtained from individual GUVs for untreated (gray), DMSO-treated (blue), and Masitinib-treated (orange) conditions at pH 7.2 and pH 4.4. An increase in the I590/I640 ratio indicates tighter lipid packing in the membrane due to Masitinib incorporation. Each data point represents an individual GUV. Statistical analysis was performed using one-way ANOVA with a confidence level of 0.05. Significance levels: *p < 0.05; ***p < 0.001; ****p < 0.0001. Scale bar: 5 μm.

GUVs were prepared using the PVA gel-swelling method and were formulated to mimic the inner leaflet of the lysosomal membrane (DOPC: DOPS: cholesterol, 40:20:40, as with Laurdan emission spectra measurement on LUVs), both at neutral pH (7.2) and acidic pH (4.4) (mol/mol, ammonium citrate, 294 mOsm/kg). Controls included GUVs in buffer (blank) and GUVs incubated with DMSO (negative control).

For both conditions tested (pH 7.2 and 4.4), GUVs did not exhibit any change in shape or morphology compared to untreated vesicles, both in the case of GUVs incubated with Masitinib or DMSO alone. Consistent with Laurdan measurements, GUVs at pH 7.2 showed no significant changes in fluorescence intensity in either channel upon incubation with DMSO or Masitinib (**Figure 4B**). Analysis of the I590/I640 ratio distribution from individual GUVs across the different treatments revealed marginal differences between untreated, drug-incubated, or DMSO-treated vesicles, with all displaying a normal distribution centered at an I590/I640 ratio of ∼1.2 (**Figure 4B**). In contrast, GUVs formed and maintained at pH 4.4 exhibited clear changes in Di-4 spectral distribution upon treatment with Masitinib (**Figure 4C**). Qualitatively, the green contribution to the GUV signal increased with Masitinib treatment, suggesting enhanced membrane packing due to drug-membrane interactions. This translated to a shift in the I590/I640 distribution toward higher values, with Masitinib-treated vesicles showing a significant population with ratios ∼2.0, which was not observed in control or DMSO-treated GUVs.

Comparison of individual spectra confirmed that at pH 7.2, Di-4 emission spectra overlapped across all conditions, with DMSO treatment only slightly reducing the 640 nm contribution (**Figure 4D**). At pH 4.4, GUVs incubated with Masitinib exhibited a pronounced decrease in the 640 nm contribution, indicating increased membrane order, while DMSO-treated and control vesicles displayed similar Di-4 emission spectra (**Figure 4E**). Collectively, the Di-4 emission spectra from individual GUVs clearly demonstrate that Masitinib induces increased lipid packing in membranes composed of DOPC: DOPS: cholesterol, and this effect occurs specifically at lysosomal-like pH (∼4.4), consistent with Laurdan spectra (**Figure 2C**). Comparison of I_590_/I_640_ distributions for all GUVs (**Figure 4F**) confirmed that while no significant changes were observed at pH 7.2 between control and Masitinib-treated vesicles, the effect at pH 4.4 was significantly stronger and statistically significant.

Overall, the changes in emission spectra of Di-4 upon interaction of Masitinib with membranes mimicking the lysosomal milieu (both in terms of lipid composition and pH) confirm the results observed with Laurdan emission spectra and further indicate that Masitinib, upon interaction with the membrane, is located at least partially in the hydrophobic region of the lipid bilayer.

### Late autophagy/lysosomal blockade enhances AKT S473 phosphorylation

In line with thermal proteome profiling data, biophysical analysis clearly indicated that masitinib accumulates within acidic membranes. Moreover, our data above also show that masitinib mediated AKT phosphorylation is sensitive to VPS34 and mTORC inhibition, suggesting a potential link to autophagy and lysosomal processes. Autophagy is an essential catabolic pathway through which cells degrade and recycle cytoplasmic components, including damaged organelles and misfolded proteins. This process involves the formation of double-membrane vesicles called autophagosomes. Upon fusion with lysosomes, autophagosomes are degraded to recycle cellular materials in the form of amino acids and metabolites, which are required for protein synthesis or ATP production [34]. Autophagy plays a critical role in cell survival under stress and in the quality control of proteins and organelles. The VPS34 and mTORC1 signaling pathways play a crucial role in regulating the initiation and progression of autophagy. The sensitivity of masitinib-induced AKT activation to these pathways raises the possibility that masitinib might affect autophagy.

To examine the impact of masitinib on autophagy, we first investigated the effect of masitinib on established markers of autophagy, specifically lipidated LC3 [35]. As shown in **Figure 5A-C**, treatment of cells with masitinib resulted in a pronounced increase in the lipidated form of LC3 (LC3-II).

**Figure 5.**
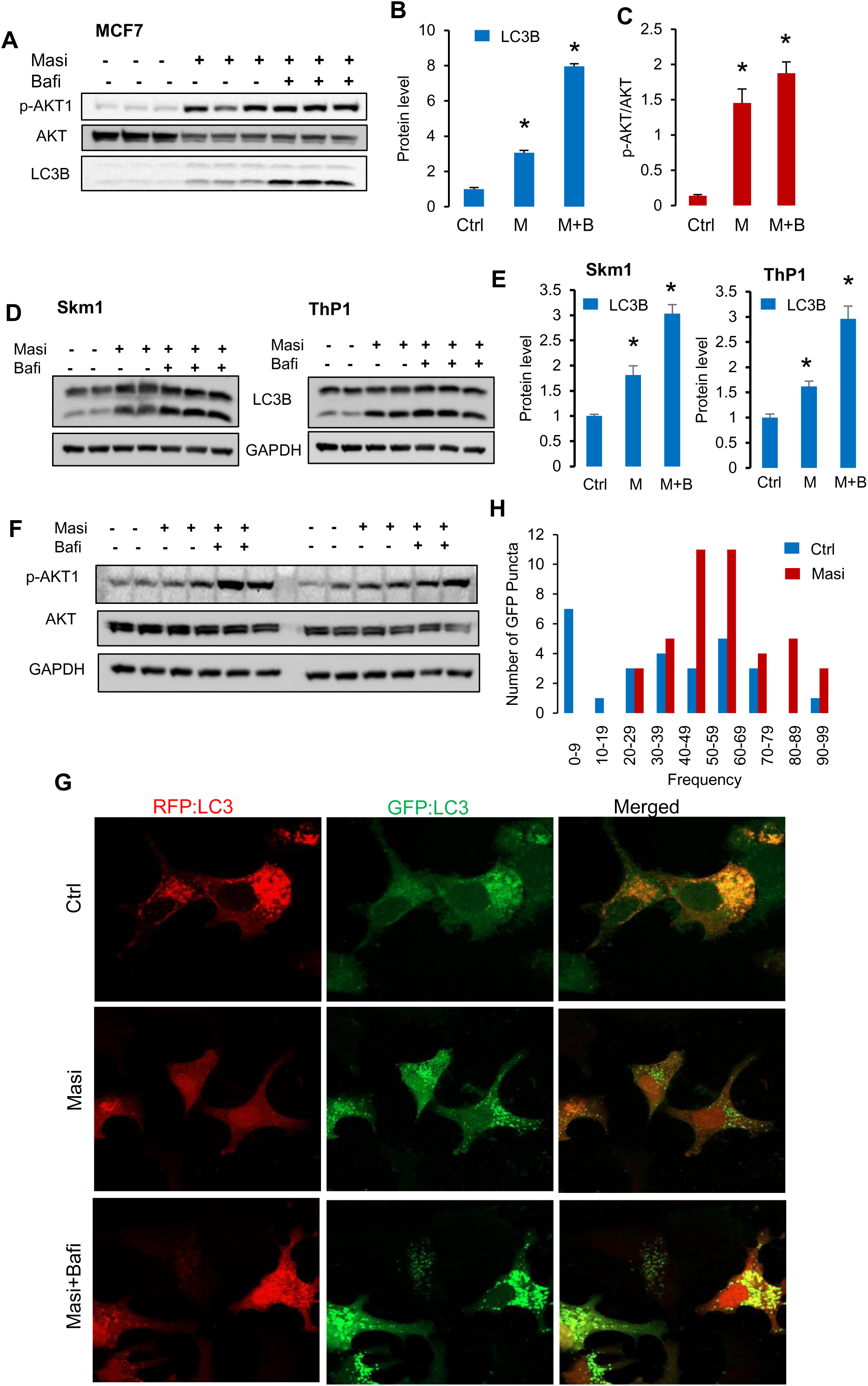
Masitinib alters autophagy flux. (**A-E**) Lysate from MCF7, Skm1, and ThP-1 cells control or treated with 5 µM Masitinib (Masi) for 24 hours, supplemented or not with 5 nM bafilomycin A for 8 hours, then stained for the indicated antibodies. **(B, C, E)** Bar graphs showing the results of densitometric analysis of the protein levels under different conditions obtained in A and D. The data are presented as the means ±SEMs. N= 3 to 4 biological replicates per condition. ANOVA, multiple comparisons: Dunnett test. **MCF7**: LC3B F (2, 6) = 813.33, P˂0.0001. p-AKT F (2, 6) = 37.65, P˂0.0005. **SKM-1**: LC3B F (2, 9) = 46.78, P˂0.0001. **THP-1**: LC3B F (2, 9) = 39.02, P˂0.0001. **(F)** Lysate from Skm1 and ThP-1 cells control or treated with 5 µM Masitinib (Masi) for 24 hours, supplemented or not with 5 nM bafilomycin A for 8 hours, then stained for the indicated antibodies. **(G)** MCF-7 cells were transiently transfected with a plasmid expressing tandem RFP-GFP-LC3 GFP-LC3. Twenty-four hours later, cells were treated with vehicle or 5 µM Masitinib. Bafilomycin A (5 nM) was added 8 hours before cell fixation. Representative image shows RFP-GFP-LC3 distribution in MCF7 cells analyzed by confocal microscopy. (**H**) Bar graph showing the frequency distribution of GFP-puncta in MCF7 control and masitinib-treated cells.

To further examine the stage at which masitinib acts within the autophagy pathway, we compared its effects with those of bafilomycin A (BafA), a well-known inhibitor of vacuolar-type ATPase that blocks autophagosome-lysosome fusion and lysosomal acidification [35]. This experiment demonstrated that co-treatment with masitinib and bafilomycin resulted in a stronger accumulation of LC3-II compared to either treatment alone (**Figure 5A-E**). This additive effect suggests that masitinib promotes autophagosome formation while simultaneously impairing autophagic flux, consistent with partial lysosomal dysfunction.

Consistent with its impact on lysosomal signaling, masitinib treatment suppressed phosphorylation of P70S6K, a downstream effector of mTORC1 [8], while enhancing phosphorylation of AKT1 (**Supplementary Figure 3A-C**). Notably, the combination of masitinib and bafilomycin further amplified AKT1 phosphorylation, indicating that lysosomal dysfunction potentiates AKT activation (**Figure 5A, 5F, Supplementary Figure 4E**). This reciprocal regulation of P70S6K and AKT1 suggests that masitinib interferes with mTORC1 activity and triggers compensatory activation of AKT through feedback mechanisms linked to lysosomal stress. Together, these findings indicate that masitinib alters lysosomal homeostasis and modulates both autophagy and mTORC-AKT signaling.

### Masitinib alters lysosome acidification and causes lysosomal damage

To investigate the effect of masitinib on the autophagy process in more detail, we used the tandem RFP-GFP-LC3 reporter to monitor autophagic flux and assess the distribution of autophagosomal puncta in control and masitinib-treated cells [36]. As shown in **Figure 4G**, masitinib treatment markedly increased the accumulation of GFP-positive puncta, indicative of impaired autophagosome-lysosome fusion or reduced lysosomal degradation. However, an increase in LC3 puncta can reflect either enhanced autophagosome formation or impaired autophagosome degradation [35]. Compared to the inhibitor alone, our data show that co-treatment of cells with BafA further increased the number of GFP-LC3 puncta (**Figure 5G-H**). This result further confirms that autophagic flux remains active and that treatment with masitinib likely promotes autophagosome formation. Finally, to examine the impact of masitinib accumulation in lysosomes and the masitinib-mediated increase in autophagosome formation on lysosomal function, we assessed lysosomal acidification using a genetically encoded, fluorescent protein-based pH biosensor. This biosensor combines a pH-sensitive mTFP1 and a relatively pH-stable mCherry, both of which are targeted specifically to the lysosome. Changes in lysosomal pH alter the mTFP1 fluorescence while mCherry serves as an internal reference, allowing ratiometric measurement of lysosomal acidification in cells [37]. As shown in **Figure 6**, masitinib-treated cells exhibited an increased number of green puncta compared to control cells, indicating a reduction in lysosomal acidification. Maintaining an acidic pH within the lysosomal lumen is essential for the activation of hydrolytic enzymes and efficient autophagic degradation. These data suggest that, while masitinib enhances autophagosome formation, its accumulation within lysosomes perturbs lysosomal activity, leading to an imbalance between autophagosome synthesis and degradation.

**Figure 6.**
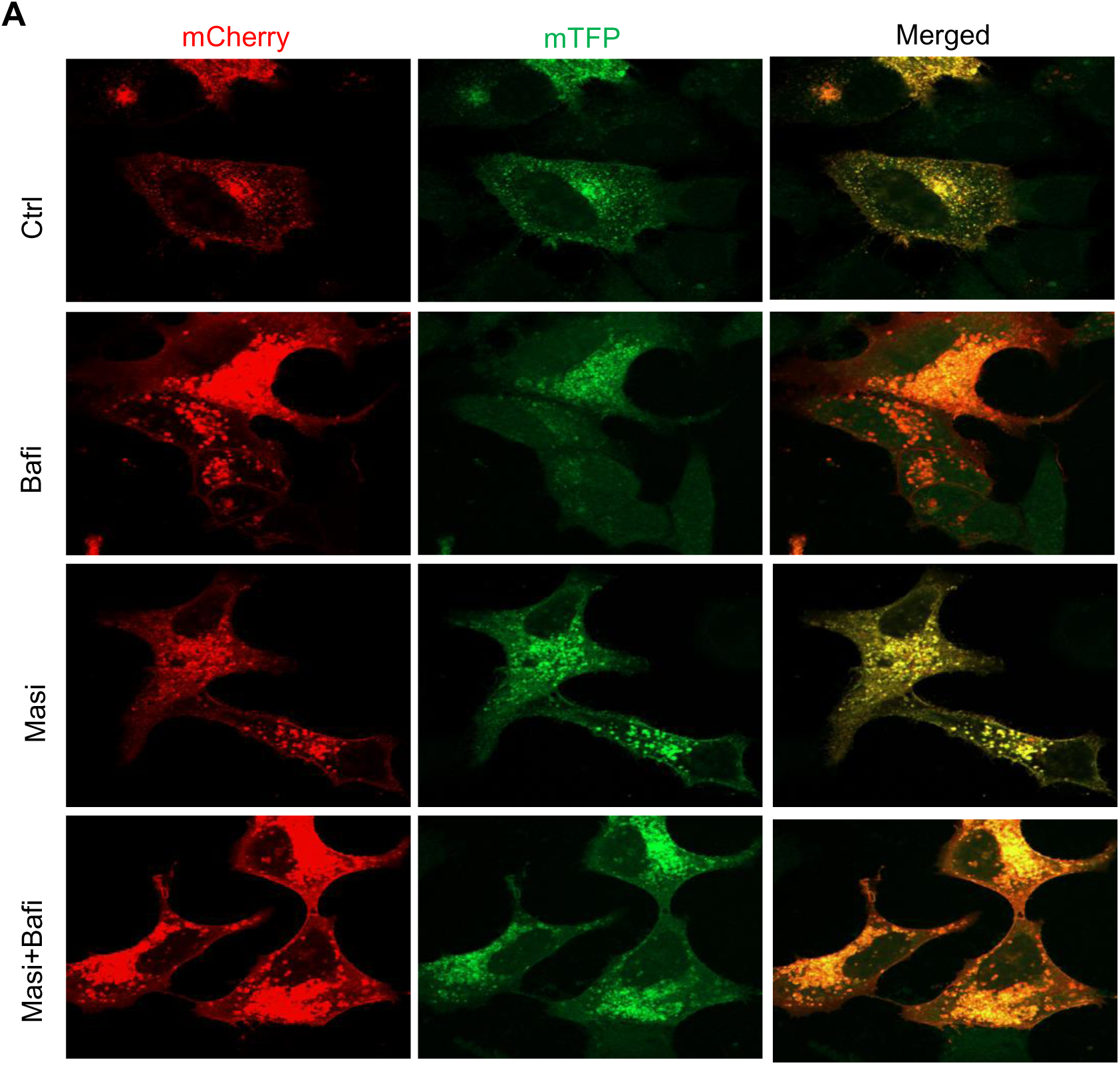
Masitinib accumulates within and affects lysosome functions. **(A)** Representative fluorescence images showing changes in lysosomal pH in MCF7 treated with vehicle or 5 μM Masitinib for 24 h, followed by treatment with bafilomycin A (5 nM) for 8 h.

## Discussion

Receptor tyrosine kinases control multiple signal transduction cascades that affect various processes, including proliferation, differentiation, and apoptosis. Moreover, RTKs play a central role in tumorigenesis, and their deregulation has been described in several cancers [7]. As a result, RTKs have become a major class for targeted therapeutics over the past years. Several RTK inhibitors have been developed and designed to target a specific receptor or a kinase domain selectively [7, 38]. One of the major challenges in the use of RTK inhibitors in cancer therapy is related to resistance mechanisms and off-target toxicities. Emerging evidence, including our data, suggests that many RTK inhibitors engage unanticipated protein targets and accumulate in specific subcellular compartments, particularly in lysosomes. Lysosomal trapping is a well-known pharmacokinetic phenomenon in which weakly basic and lipophilic drugs become sequestered in the acidic environment of lysosomes. This accumulation not only alters both the intracellular distribution of the drugs but also their functional activity, which often diverts them from their intended targets and causes them to interact with lysosomal proteins instead [11–12, 39]. Among these inhibitors is Masitinib, initially used for the treatment of mast cell tumors in animals [1], and recently tested in clinical trials to target mast cells and microglia for the treatment of neurological and inflammatory diseases, including amyotrophic lateral sclerosis, multiple sclerosis, and Alzheimer’s disease [3–6]. However, due to concerns related to the reliability of the study results, the European Medicines Agency refused marketing authorization for Masitinib [40]. Using thermal proteome profiling, our data demonstrated that lysosomal proteins as the most thermally altered upon masitinib treatment. In line with our observations, Neuwirt et al. [14] recently reported that several receptor tyrosine kinase inhibitors, including masitinib, induce lysosomal damage and cathepsin-dependent destabilization of lysosomal membranes. Together, these findings suggest that lysosomes represent a previously underappreciated cellular compartment affected by masitinib.

Our thermal proteomics data revealed marked changes in the thermal stability of lysosomal proteins. This effect could be linked to strong accumulation of masitinib within lysosomes and their membranes, which in turn alters the thermal stability of resident lysosomal proteins. To further examine whether lysosomal accumulation is the main driver of the observed thermal shifts, we performed time-resolved Laurdan fluorescence assays using large unilamellar vesicles enriched in negatively charged lysosomal lipids [17, 30]. These experiments confirmed that masitinib preferentially interacts with membrane lipids, mimicking the physicochemical properties of lysosomal membranes (namely, the presence of negatively charged lipids), particularly under acidic conditions that resemble the lysosomal milieu. Increasing the compositional complexity of the liposomes to further mirror the lysosomal membrane maintains the shift in Laurdan GP under acidic conditions. These results highlight how the combination of negatively charged lipids at the membrane surface and low pH is the driving force for Masitinib-membrane interaction.

Similarly, by complementing measurements of Laurdan emission with Di-4 spectral analysis on GUVs, we confirmed that both the presence of negatively charged lipids and acidic pH (i.e., lysosomal milieu) are required for Masitinib intercalation within membranes. Since Di-4 fluorescence emission shift is sensitive to the presence of molecules within the hydrophobic region of the lipid bilayer, we can infer that Masitinib interaction at the membrane might not be a simple adsorption at the surface but a bona fide insertion within the membrane.

Our data show that masitinib treatment led to a marked inhibition of mTORC1 signaling, as shown by reduced phosphorylation of P70S6K. This inhibition could be directly linked to the accumulation of masitinib in lysosomes, which disrupts basal autophagy and lysosomal function, which in turn leads to lower intracellular amino acid and ATP availability, thereby suppressing mTORC1 [41]. Consistent with this hypothesis, treatment of cells with masitinib leads to a marked increase in LC3 lipidation and impaired lysosomal acidification.

Unexpectedly, masitinib also induced a strong increase in AKT phosphorylation, which was further enhanced by bafilomycin treatment, suggesting that AKT activation in this case could be linked to activation of a stress response pathway. Our data show that mTORC2 contributed only moderately to this effect, because Rictor depletion did not completely block AKT phosphorylation. This indicates that AKT activation occurs largely through a non-canonical pathway. In this respect, several kinases have been shown to phosphorylate AKT independently of mTORC2, including DNA-PK in response to DNA damage and TBK1, which localizes to endolysosomal membranes and integrates stress and nutrient-sensing pathways [18]. It is therefore possible to imagine that TBK1 or other related kinases mediate the paradoxical activation of AKT observed upon masitinib treatment.

Preclinical studies in rodents have suggested that masitinib exhibits promising neuroprotective properties by reducing neuroinflammation and pro-inflammatory cytokines [3–4]. However, clinical findings in humans have been inconsistent, and in some cases, have revealed adverse effects [5–6]. The reasons behind these discrepancies are unknown; however, our findings provide a potential mechanistic explanation. The anti-inflammatory effects seen upon short-term treatment in preclinical models may be beneficial, but sustained lysosomal stress and autophagy impairment triggered by masitinib could compromise its long-term therapeutic efficacy in humans.

Together, these findings highlight the importance of further exploring the cellular consequences of lysosomal dysfunction induced by masitinib. Indeed, it will be interesting to determine how lysosomal trapping of masitinib influences tumor cell growth and cell viability in cancer models, especially in solid tumors that are highly dependent on autophagy for survival. Moreover, future work should also address whether preventing lysosomal accumulation of masitinib could lower cellular stress responses and improve its therapeutic potential.

## Materials and Methods

### Reagents

RPMI1640 with L-Glutamine (Capricorn Scientific, #RPMI-A), DMEM High Glucose (4.5 g/l), with L-Glutamine, Sodium Pyruvate (Capricorn Scientific, #DMEM-HPA), Fetal Bovine Serum (Sigma Aldrich, # F7524-500ML), Penicillin-Streptomycin (Thermo Fisher Scientific, #15140122). The following inhibitors were used: Masitinib (Tocris Bioscience, #7705), Bafilomycin A1 (Sigma Aldrich, #B1793), GDC-0980 (MOLNOVA, #M10158), Rapamycin (Thermo Scientific Chemicals, #J62473.MC).

Chloroform solutions of POPC (1-palmitoyl-2-oleoyl-glycero-3-phosphocholine, Mw= 760.076), DOPC (1,2-dioleoyl-sn-glycero-3-phosphocholine, Mw: 786.11), DOPE (1,2-dioleoyl-sn-glycero-3-phosphoethanolamine, Mw: 744.034), DOPS (1,2-dioleoyl-sn-glycero-3-phospho-L-serine, Mw: 810.025), 18:1 BMP S, S (sn-(3-oleoyl-2-hydroxy)-glycerol-1-phospho-sn-1’-(3’-oleoyl-2’-hydroxy)-glycerol, Mw: 792.075), and Egg Sphingomyelin were purchased from Avanti Polar Lipids. Di-4-ANEPPDHQ was provided by ThermoFisher Scientific.

Sucrose, Glucose, sheep wool cholesterol (Mw:386.65), and Laurdan (6-dodecanoyl-N, N-dimethyl-2-naphthylamine) were purchased from Sigma-Aldrich.

### Cell culture

MCF7 cells were generously provided as a gift from Dr. Anna Marusiak, IMol Institute. The cells were cultured in DMEM supplemented with 10% fetal bovine serum (FBS), 100 U/mL penicillin, and 100 U/mL streptomycin. SKM1, KG-1, Molm14, and THP-1 cell lines gift from (Gift from Dr. Natalia Baran, Instytut Hematologii i Transfuzjologii, Warsaw) were grown in RPMI1640 medium supplemented with 10% fetal bovine serum (FBS), 100 U/mL penicillin, and 100 U/mL streptomycin. Cells were maintained at 37°C in a humidified 5% CO2 incubator. The concentrations and durations of each treatment of the cell with the inhibitors listed above were all described in the figure legends. Experiment control cells were treated with an equivalent volume of solvent (DMSO).

### Immunofluorescence

MCF7 cells were first plated overnight in 12-well plates on glass coverslips (50000 cells/well). Adherent cells were then transfected with the following plasmids: pcDNA3-GFP-LC3-RFP-LC3ΔG was a gift from Noboru Mizushima (Addgene plasmid # 168997; http://n2t.net/addgene:168997; RRID: Addgene_168997), and pLJM1-FIRE-pHLy was a gift from Aimee Kao (Addgene plasmid # 170775; http://n2t.net/addgene:170775; RRID: Addgene_170775). Transfection was performed using JetPRIME transfection reagent (Polyplus, #101000027) according to the manufacturer’s instructions. Twenty-four hours later, cells were treated with vehicle or 5 µM Masitinib for 24 hours, followed by 8 hours of 5 nM Bafilomycin A. The cells were then fixed using 4% formaldehyde. After a series of three washes with PBS, Glass coverslips were then mounted on slides with mounting solution without DAPI (Invitrogen #P36961). Images were captured using a Zeiss LSM 710 confocal microscope and processed with ImageJ (National Institutes of Health, Bethesda, MD, USA).

### Proteome Integral Solubility Alteration Analysis of Masitinib

PISA assay was carried out as described previously [19]. Briefly, SKM1 cells were seeded into a 75 cm culture flask. Thirty minutes later, cells were treated for two hours with 20 μM Masitinib or DMSO as a control, with three biological replicates for each condition. After treatment, cells were collected by centrifugation (2500 rpm, 5min) and washed twice with phosphate-buffered saline (PBS). Cell pellets were collected and resuspended in PBS. After homogenization, the content of each biological replicate was divided into eight 0.2 mL PCR tubes, with 60 μL per tube, one for each temperature point. The following gradient temperatures were used for PISA (43°C, 44.3°C, 46.2°C, 48.5°C, 51.8°C, 54.3°C, 55.9°C, 57°C). The cells were treated for 3 minutes with the indicated temperatures using a C1000 Touch Thermal Cycler (BioRad), followed by 5 minutes of incubation at room temperature.

Cell lysates were isolated as follows: first, three freeze-thaw cycles were performed using liquid nitrogen, and then the samples were thawed at 30 °C. Next, aliquots of the same biological replicates were recombined and incubated for 15 minutes in 0.2% NP-40 in PBS at 4°C using a thermal shaker under constant agitation. Soluble proteins were obtained after two rounds of centrifugation (20 min, 20,000 g, 4 °C).

Protein levels were quantified using the Pierce™ BCA Protein Assay Kit (Thermo Scientific™, #23225). For each biological replicate, 20 μg of total protein was subjected to overnight trypsin digestion at 37 °C with agitation at 1000 rpm. Following digestion, samples were processed for TMT labeling. Peptide cleanup columns were prepared by packing three layers of C18 resin mesh into 200 µL pipette tips to create STAGE tip columns, which were then mounted onto 1.5 mL microcentrifuge tubes. The columns were conditioned by centrifugation with 150 µL of methanol (1200 g, 2 min), followed by a wash with 100 µL of 50% acetonitrile containing 0.1% formic acid (1200 g, 2 min). The resin was then equilibrated twice using 150 µL of 0.1% formic acid using the same centrifugation conditions. Digested peptides corresponding to 10 µg were loaded onto each C18 STAGE tip and centrifuged (1200 g, 2 min), after which the columns were washed twice with 150 µL of 0.1% formic acid. TMTs labels were first dissolved in 2 µL acetonitrile and diluted with 200 µL of freshly prepared 50 mM HEPES (pH 8). Peptide labeling was carried out by loading 200 µL of TMT mixture onto the columns and centrifuged at 300 g for 10 minutes. Upon three washes with 150 µL of 0.1% formic acid (1200 g, 2 min), labeled peptides were eluted using 60 µL of 60% acetonitrile and collected after centrifugation (1200 g, 2 min). A volume of 55 µL was taken from each sample and combined, then dried in a centrifugal vacuum concentrator at 40 °C (SpeedVac). Samples were then reconstituted in 0.1% trifluoroacetic acid and fractionated with the Pierce High pH Reversed-Phase Peptide Fractionation Kit (Thermo Scientific™, #84868) following the manufacturer’s procedure. Fractions were again dried by SpeedVac at 40 °C, resuspended in 0.1% formic acid, and subsequently analyzed by LC-MS/MS.

### LC-MS/MS measurements and data analysis

Mass spectrometry analyses were carried out at the Mass Spectrometry Laboratory, Institute of Biochemistry and Biophysics PAS. Each fraction was dissolved in 100 µl of solvent A (0.1% formic acid in water) using 30 min of sonication and subsequently mixed on a vortex for 30 min. A volume of 45 µl of each fraction was injected into the LC-MS system. Peptide analysis was performed using an Evosep One system (Evosep Biosystems) coupled online to an Orbitrap Exploris 480 mass spectrometer (Thermo Fisher Scientific). Sample loading onto Evotips Pure C18 (Evosep Biosystems) was carried out according to the manufacturer’s guidelines. Liquid chromatography was conducted at 500 nl/min using the 88-min predefined method on an EV1137 analytical column (Dr Maisch C18 AQ, 1.5 µm particles, 150 µm ID, 15 cm, Bruker). Data were acquired in positive-ion mode using a data-dependent acquisition workflow. Full MS scans (MS1) were recorded at a resolution of 60,000 with a normalized AGC target of 300%, Auto injection time, and a mass range of 300-1700 m/z. MS/MS scans (MS2) were collected at a resolution of 30,000 with Standard AGC settings, Auto injection time, and selection of the 25 most intense precursors using a 1.2 m/z isolation window. A dynamic exclusion period of 20 s was applied with a ±10 ppm mass tolerance. The minimum precursor signal threshold was set at 5×10³, and the Precursor Fit filter was enabled at 70%. Fragmentation was performed with a normalized collision energy of 30%. TurboTMT mode was configured for TMTpro reagents. Source parameters included a spray voltage of 2.1 kV, a funnel RF level of 40, and a capillary temperature of 275 °C.

The data were processed with MaxQuant v. 2.4.3.0, and the peptides were identified from the MS/MS spectra searched against the Uniprot reference human proteome (UP000005640) using the build-in Andromeda search engine. Cysteine carbamidomethylation was set as a fixed modification and methionine oxidation, glutamine/asparagine deamidation, and protein N-terminal acetylation were set as variable modifications. For in silico digests of the reference proteome, cleavages of arginine or lysine followed by any amino acid were allowed (trypsin/P), and up to two missed cleavages were allowed. FAIMS-Reporter ion MS2-based quantification was applied with reporter mass tolerance = 0.003 Da and min. reporter PIF = 0.75. The FDR was set to 0.01 for peptides, proteins and sites. Match between runs was enabled. Other parameters were used as preset in the software. Unique and razor peptides were used for quantification enabling protein grouping (razor peptides are the peptides uniquely assigned to protein groups and not to individual proteins). Reporter intensity corrected values were uploaded into Perseus v. 1.6.10.0 [42]. Standard filtering steps were applied to clean up the dataset: reverse (matched to decoy database), only identified by site, and potential contaminants (from a list of commonly occurring contaminants included in MaxQuant) protein groups were removed. The intensity values were log2 transformed and protein groups with all values were kept. The values were then normalized by median subtraction within TMT channels. Proteins exhibiting masitinib-induced changes in thermal stability were identified by statistical analysis. Two-tailed Student’s t-tests (n=3, p<0.01) were performed for protein groups quantified across masitinib and DMSO conditions. Results were then filtered to retain only protein groups supported by more than one razor+unique peptide and showing an absolute log2 fold change greater than 0.3, yielding a high-confidence set of proteins with significantly altered thermal stability upon masitinib treatment. This dataset has been deposited to the ProteomeXchange Consortium via the PRIDE [43–44] partner repository with the dataset identifier PXD071058.

#### Western Blotting

Cells treated with vehicle or the indicated inhibitors were washed once with PBS and then lysed in a buffer containing 50 mM Tris-HCl (pH 8.0), 150 mM NaCl, 1.0% NP-40, 0.1% SDS, 25 U/mL Benzonase, and 100 µM PMSF (Thermo Scientific™, #36978). Cell lysates were cleared by centrifugation at 12,000 × g for 10 minutes at 4°C, and protein concentrations were quantified using a BCA assay (Thermo Scientific™, #23225). Equal amounts of protein (15-25 µg) were separated by SDS-PAGE and transferred onto nitrocellulose membranes. Following transfer, membranes were blocked for 1 hour in TBST (0.05% Tween-20) containing 5% non-fat milk and incubated overnight at 4°C with primary antibodies. After three TBST washes, membranes were incubated with HRP-conjugated secondary antibodies for 1 hour at room temperature, and signals were detected using the Amersham ImageQuant 800 system.

Primary antibodies were obtained from Invitrogen™: phospho-AKT1 (Ser473) (#700256), AKT Pan (#MA5–14916), LC3B (#PA1-46286), Phospho-p70 S6 Kinase (Thr389) #710095, P70 S6 Kinase #MA5-15141 and GAPDH (#MA5-15738). Secondary antibodies were obtained from Invitrogen™: Goat anti-Mouse IgG (H+L) Secondary Antibody, HRP (#31430) and Goat anti-Rabbit IgG (H+L) Secondary Antibody, HRP (#31460).

#### Cell growth assay

To assess the effect of masitinib on cell growth, 20,000 cells per well of the indicated cell line were seeded in 96-well plates and treated with the indicated concentrations of masitinib or vehicle control. After 72 hours, the metabolic activity of the cells was evaluated using the MTT assay. This colorimetric assay is based on the reduction of 3-(4,5-dimethylthiazol-2-yl)-2,5-diphenyltetrazolium bromide (MTT) to insoluble formazan crystals by metabolically active cells, serving as an indicator of cell viability. A volume of 10 μL of MTT labeling reagent (final concentration 0.5 mg/mL) was added to each well, and the plates were incubated for 4 hours at 37°C in a humidified atmosphere containing 5% CO₂. Following incubation, 100 μL of SDS-HCl solution was added to each well to solubilize the formazan crystals. The plates were incubated for an additional 4 hours under the same conditions. Samples were then mixed by pipetting, and the absorbance was measured at 570 nm using a microplate reader (Multiskan™ FC Microplate Photometer, Thermo Scientific™).

### Large unilamellar vesicles preparation

Large unilamellar vesicles (LUVs) were prepared following established methods [45]. Briefly, lipid solutions in chloroform, containing various phospholipid species, were mixed at desired molar ratios in a glass vial. The organic solvent was evaporated under vacuum for 12 hours. Prior to evaporation, 1 mol% Laurdan in chloroform was added to the lipid solution. The resulting lipid film was hydrated with either phosphate-buffered saline (PBS, pH 7.2) or ammonium citrate (pH 4.4) at 40°C to achieve the target concentration, followed by gentle vortexing. The multilamellar vesicle suspension was sonicated for 10 minutes to break up larger aggregates and then extruded 21 times through 100 nm pore-size polycarbonate filters (Avanti Polar Lipids) using a mini-extruder (Avanti Polar Lipids). Vesicle size and concentration were confirmed by dynamic light scattering (DLS), and the liposomal suspension was used within two weeks of extrusion. All chemicals were of high purity (>99%) and used without further purification.LUVs were formulated to mimic specific membrane compositions: (1) early endosomal membranes (DOPC: cholesterol, 60:40 mol/mol, PBS, pH 7.2, 297 mOsm/kg) [30]; (2) the inner leaflet of lysosomal membranes, enriched in negatively charged lipids (DOPC:DOPS: cholesterol, 40:20:40 mol/mol, ammonium citrate, pH 4.4, 294 mOsm/kg) [31]; or (3) the complex lipid composition of lysosomal membranes (POPC:DOPE:sphingomyelin:BMP: cholesterol, 25:5:30:15:25 mol/mol, in either PBS, pH 7.2, 297 mOsm/kg, or ammonium citrate, pH 4.4, 294 mOsm/kg) [33].

### Giant unilamellar vesicles preparation

GUVs were prepared via gel-assisted spontaneous swelling as previously described [46]. Briefly, a 5% (w/w) PVA solution was prepared by stirring PVA in PBS (297 mosm/kg) while heating at 90°C. PVA-coated substrates were prepared by spreading 100–300 μL of the PVA solution onto a frosted 30 mm diameter microscope coverslip (Menzel-Gläser), followed by drying for 30 minutes at 50°C. Prior to use, coverslips were cleaned with UV/Ozone for 15 minutes to prevent dewetting of the PVA film. Subsequently, 10-20 μL of lipids (DOPC: DOPS: cholesterol, 40:20:40 mol/mol) dissolved in chloroform (1 mg/mL) were spread over the dried PVA film and placed under vacuum for 30 minutes to remove solvent.

A chamber was assembled by placing a silicone hole gasket on the PVA film to center the lipid layer. The chamber was preheated to 50°C on a heating block, then filled with 300 µL of the appropriate buffer-either 50 mM sucrose in PBS (pH 7.2, 330 mosm/kg) or 50 mM sucrose in ammonium citrate (pH 4.4, 340 mosm/kg)-preheated to 50°C. The chamber was sealed with a second glass coverslip.

After typically 2 hours at 50°C, once GUVs reached the desired size, they were gently collected into an Eppendorf tube using a pipette. The vesicle suspension was diluted 5-fold with 50 mM glucose in either PBS (pH 7.2) or ammonium citrate (pH 4.4) and incubated undisturbed for 1 hour before use. GUVs were then stained with ∼0.5% Di-4 post-formation via incubation at 37°C for 30 min.

### Laurdan fluorescence spectra kinetics

Laurdan fluorescence spectra were measured using a Synergy Neo plate reader on a 96-well black plate with an opaque bottom. For each well, 150 µL of liposomal suspension at 150 µM was added, followed by Masitinib in DMSO to achieve a 1% volume fraction (final Masitinib concentration: 5 µM). Control wells included LUVs in buffer only (blank) and LUVs with 1% DMSO alone (negative control). Samples were incubated at 37°C for 8 hours, and Laurdan GP was monitored by measuring fluorescence intensity every 10 minutes with orbital shaking. Emission was recorded at 440 nm and 490 nm (slit width: 10 nm, gain: 100, integration time: 20 ms) upon excitation at 350 nm (slit width: 5 nm).

Laurdan GP kinetics curves were calculated using the General Polarization equation [16]: GP = (I440 – I490)/(I440+I490) Where I440 and I490 are the fluorescence intensity measured at 440 nm and 490 nm, respectively. For each condition, we acquired three (n=3) technical replicates.

### Di-4-ANEDDPHQ confocal spectral imaging

Di-4 labeled GUVs suspension at 150 uM lipid concentration was placed on a 35mm imaging dish, mixed with 1% DMSO (vol/vol) or Masitinib in DMSO at 1% (v/v) to reach a final concentration of 5 µM Masitinib, and let sit for 20 minutes to allow GUV sedimentation via differential density between luminal sucrose and bulk glucose.

GUVs were visualized with a ZEISS LSM 900 confocal microscope, using a 60x oil-immersion objective, and the GUVs signal was acquired in two distinct modalities:

i) Fluorescence confocal mode with excitation wavelength of 470 nm using two defined emission channels corresponding to the green (505 – 600 nm) emission window and red (605 – 650 nm) emission window of the Di-4 spectra, respectively.
ii) Di-4 spectral imaging using the lambda stack modality, with excitation 470 nm, emission 505–650 nm range, split into 16 channels of 8 nm each, to derive the full emission spectra of Di-4 from the images.

For each condition and acquisition mode, we acquired at least three 1 mm × 1 mm fields of view (FOV) via tile stitching combined with z-stacking (10 µm z-width, 0.5 µm z-step) to capture the equatorial projection of all GUVs present in the field of view.

Following acquisition, each GUVs was cropped from the FOV, and the corresponding I590/I640 and Di-4 spectra were extrapolated from spectral image stacks and exported using the ZEISS Zen software. Subsequent analysis of spectra distributions between conditions was performed using OriginPro software.

#### Statistical analysis

Statistical analysis was performed using GraphPad Prism (http://www.graphpad.com). All experiments were carried out using at least three biological replicates. Comparisons between two groups were analyzed using an unpaired, two-tailed t-test. For comparisons among three or more groups, a one-way ANOVA followed by Dunnett’s post hoc test was employed. Differences were considered statistically significant when P < 0.05.

## DECLARATIONS

The AI Grammarly was used at the final stage of the manuscript preparation for English corrections and clarifications.

### Ethics approval and consent to participate

not applicable.

### Consent for publication

not applicable.

### Availability of data and materials

The datasets used and/or analysed during the current study are available from the corresponding author on reasonable request.

### Authors’ contributions

Conceptualization, A.A. and M.K., Methodology; A.A., M.K., and R.S, Proteomic sample preparation and data analysis: A.A., R.S; Investigation, A.A., M.K., and A.E.S.; original draft preparation, AA, M.K.; Writing review and editing, A.A., M.K., and R.S.; Funding acquisition, A.A.

## Acknowledgment

The authors thank the IMol Proteomics Core Facility for proteomic sample preparation and the Mass Spectrometry Laboratory, Institute of Biochemistry and Biophysics PAS, for PISA sample analysis.

## Supplementary Figures

**Supplementary Figure 1.**
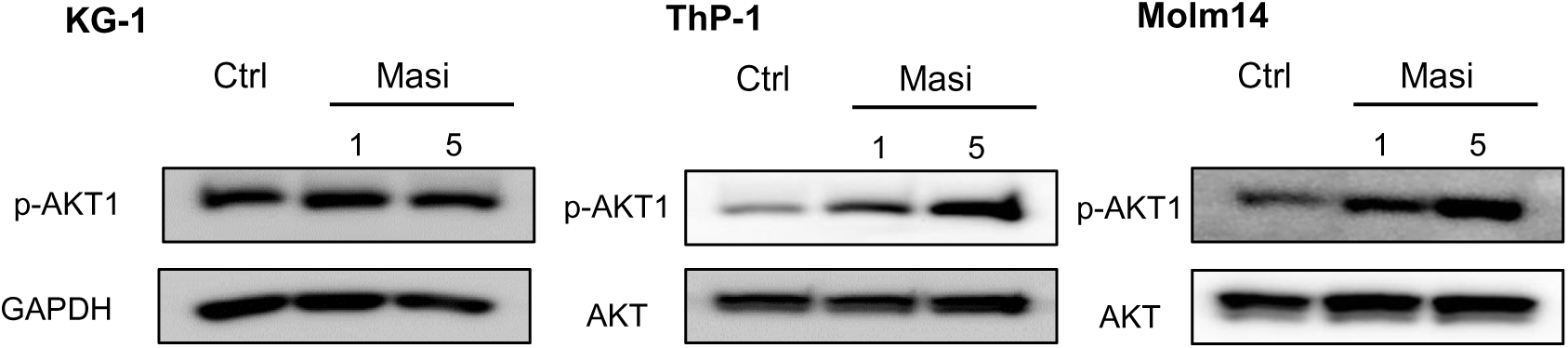
(A) Lysate from KG-1, ThP-1, and Molm14 cells control or treated with the indicated concentrations of mastinib (µM) for 24 hours, then stained for the indicated antibodies.

**Supplementary Figure 2.**
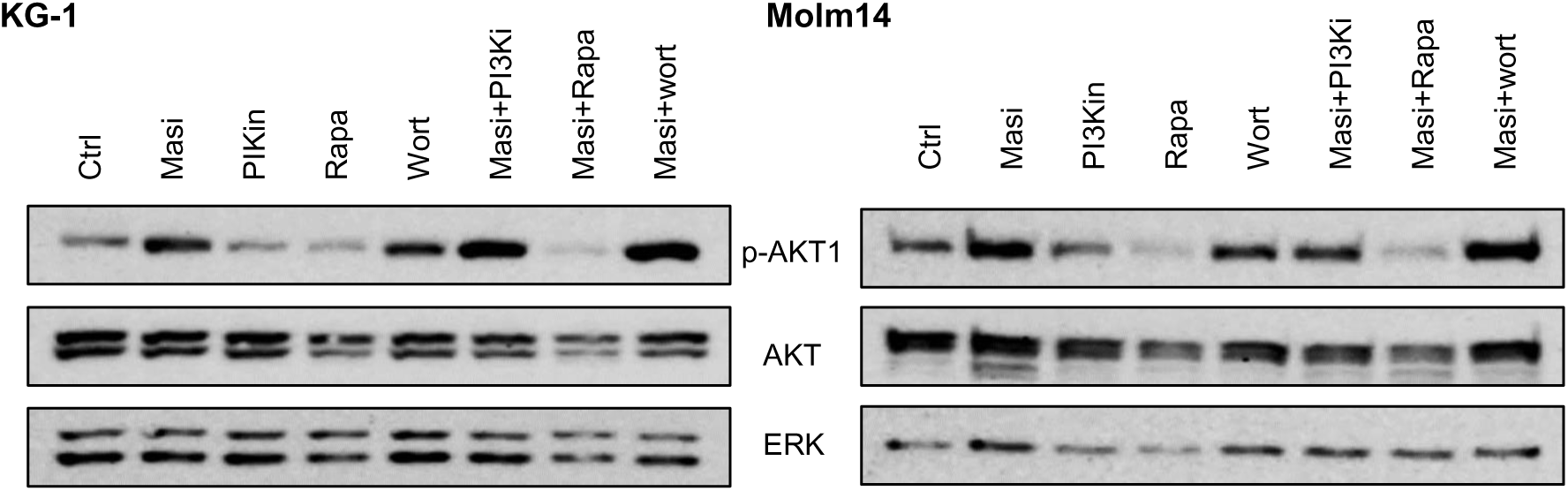
(A) Lysate from KG-1 and Molm14 cells control or treated with 5 µM Masitinib (Masi) supplemented with the indicated inhibitor for 24hours, then stained for the indicated antibodies. PI3Ki (GDC-0980, 50nM), Rapamycin 50nM, wortmannin 50nM.

**Supplementary Figure 3.**
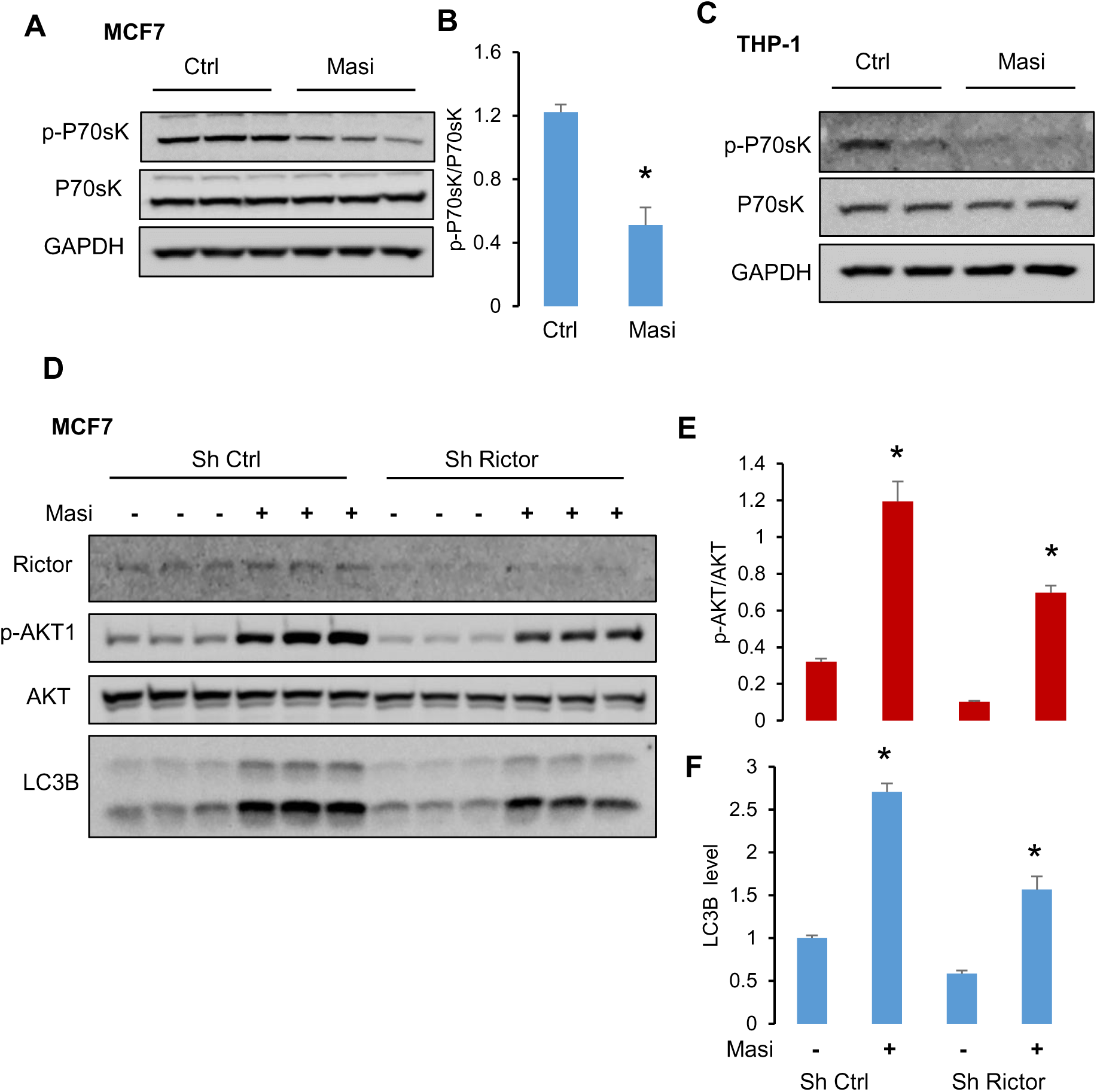
(A, C) Lysate from MCF7 and THP-1 cells control or treated with 5 µM Masitinib (Masi) for 24 hours, then stained for the indicated antibodies. (B) Bar graphs showing the results of densitometric analysis of the protein levels under different conditions obtained in A. Unpaired T-test p<0.005. (D) Lysate from MCF7 cells stably expressing sh control or sh against Rictor treated or not with 5 µM Masitinib for 24 hours, then stained for the indicated antibodies. (E-F) Bar graphs showing the results of densitometric analysis of the protein levels under different conditions obtained in D. The data are presented as the means ±SEMs. ANOVA, multiple comparisons: Dunnett test. p-AKT1 F (3, 8) = 52.39, P˂0.005, LC3B F (3, 8) = 96.92, P˂0.0001.

**Supplementary Figure 4.**
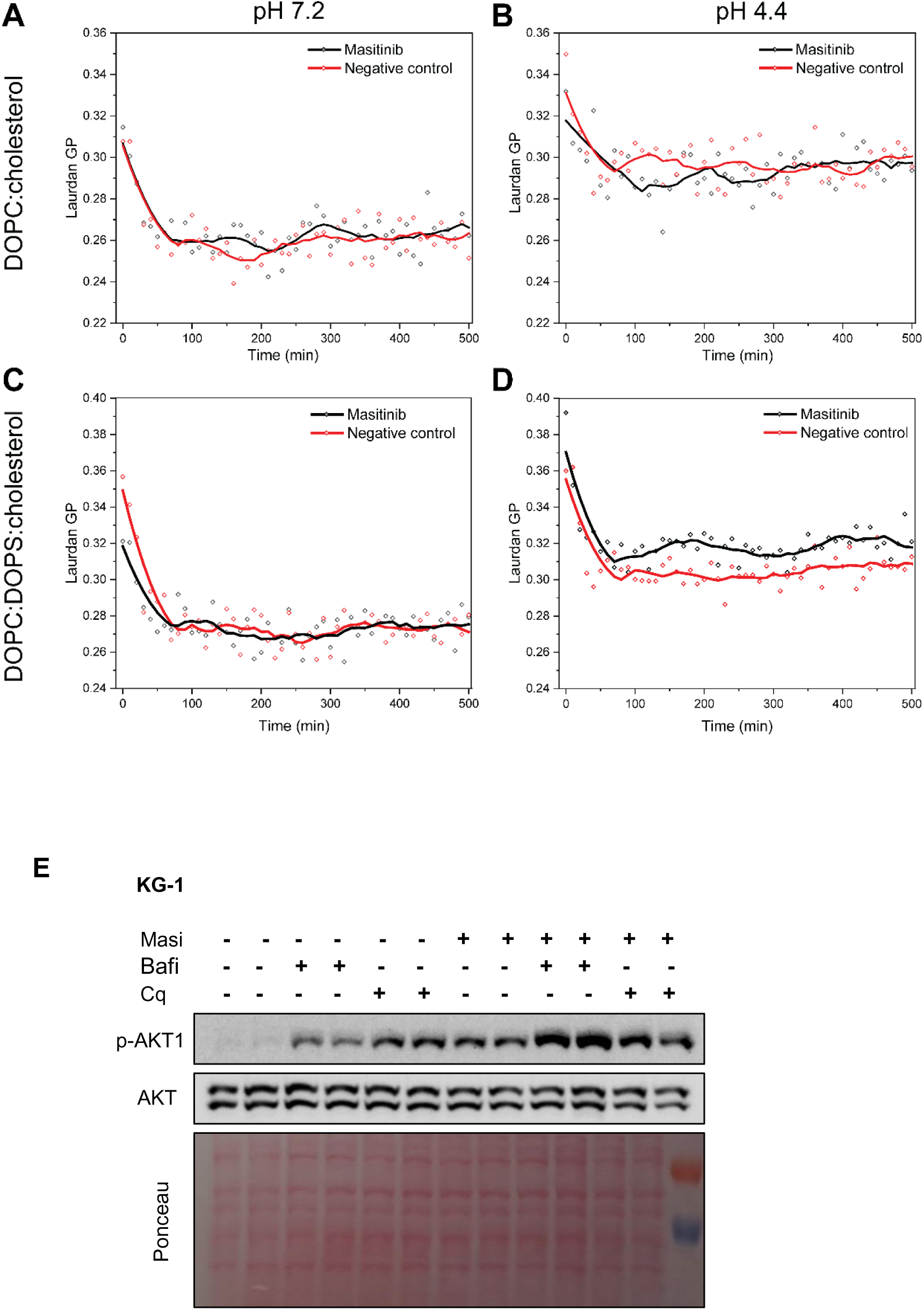
Representative kinetics profile of Laurdan GP for liposomes composed of either DOPC: cholesterol 60:40 (A-B) and DOPC:DOPS: cholesterol 40:20:40 (C-D) at pH 7.2 and pH 4.4, incubated with 5 μM Masitinib in DMSO (black) or DMSO only (negative control, red). Data symbols represent each GP measurement, while the line represents the kinetic profile obtained from smoothing the scatter plot using a 15-point average. (E) Lysate from KG-1 cells control or treated with 5 µM Masitinib (Masi) supplemented or not with 5 nM bafilomycin A or 100nM chloroquine for 24 hours, then stained for the indicated antibodies.

